# An FGF Timer for Zygotic Genome Activation

**DOI:** 10.1101/2022.10.01.510450

**Authors:** Nicholas Treen, Emily Chavarria, Claire J. Weaver, Clifford P. Brangwynne, Michael Levine

**Author notes:** Corresponding author - Nicholas Treen.

## Abstract

Zygotic genome activation has been extensively studied in a variety of systems including flies, frogs, and mammals. However, there is comparatively little known about the precise timings of gene induction during the earliest phases of embryogenesis. Here we employ high-resolution *in situ* detection methods, along with genetic and experimental manipulations, to study the timing of zygotic activation in the simple model chordate, *Ciona intestinalis* with minute-scale temporal precision. We found that two Prdm1 homologs in *Ciona* are the earliest genes that respond to FGF signaling. We present evidence for a FGF timing mechanism that is driven by derepression of the ERF repressor by ERK activity, which works in concert with localized activators such as *Foxa*.*a*. Absence of ERF results in derepression of target genes throughout the embryo. A highlight of this timer is the sharp transition in FGF responsiveness between the 8- and 16-cell stages of development. We propose that this timer is an innovation of chordates that is also employed by other vertebrates.

## Introduction

The earliest stages of embryonic development are remarkable, as the cells are not transcribing RNA and function purely through maternal contributions. This abruptly changes at precise points in development when the zygotic genome begins to transcribe mRNA and maternal mRNAs are depleted (Vastenhouw et al. 2019).

In the ascidian *Ciona*, zygotic genome activation occurs between the 8- to 32-cell stages (Lamy et al. 2006; Satou 2020). At the 8-cell stage, transcription is extremely limited, with *Foxa*.*a* being one of the only genes expressed (Lamy et al. 2006; Treen et al. 2018). At the 16- cell stage a greater set of genes are expressed and there is some lineage specification (Oda-Ishii et al. 2016; Treen et al. 2018). By the 32-cell stage there is full zygotic genome activation with the germ layers being mostly specified (Satou et al. 2020; Tokuoka et al. 2021). Additionally, at the 32-cell stage there is a sufficiently complicated embryonic geometry, as well as activating and inhibiting signals, for neural induction to occur through FGF/ERK signaling resulting in the expression of *Otx* in the a6.5 and b6.5 blastomeres (Bertrand et al. 2003; Hudson et al. 2003; Ohta and Satou 2013; Williaume et al., 2021).

Genes activated by cell signaling pathways are generally repressed in the absence of an inductive signal (Barolo and Posakony 2002; Affolter et al. 2008). For example, the Ets-class transcriptional repressor ERF is known to repress target genes in the absence of ERK phosphorylation. Activation of ERK by RTK signaling pathways such as FGF triggers ERF phosphorylation and derepression of its target genes (Sgouras et al. 1995; Le Gallic et al. 1999).

We previously determined that ERF undergoes phase separation in response to FGF signaling. When FGF is active, ERF is phosphorylated, causing condensates to dissolve, coinciding with derepression of target genes (Weaver et al. 2022). ERF was also shown to repress *Otx* in *Ciona* embryos at the 32-cell stage (Williaume et al. 2021). Interestingly, we previously observed that ERF condensates were able to dissolve and reform within a single interphase (Weaver et al. 2022), suggesting a dynamic response to endogenous FGF signals.

We identified *Prdm1-r*.*a* and *Prdm1-r*.*b* as the earliest target genes that are regulated by FGF signaling at the 16-cell stage. However, unlike *Otx*, this activation does not depend on cell-cell contacts. Here we show that three FGF responsive genes, *Prdm1-r*.*a, Prdm1-r*.*b* and *Otx*, are activated in a precise temporal order from the 8- to 32- cell stages. Sequential expression depends on derepression of ERF. Perturbing ERF results in derepression of all three genes at the 16-cell stage. We show that expression of *Prdm1-r*.*a/b* occurs due to competition between the repressor ERF and the activator Foxa.a. Altogether, these results suggest that a FGF timer acts through derepression of ERF at the 16-cell stage. This timer is then augmented by high levels of FGF signals at the 32-cell stage. The consequence of these processes is the precise temporal activation of the earliest genes during zygotic genome activation.

## Results and Discussion

*ERF* is present as a maternal mRNA in *Ciona* unfertilized eggs and early embryos (Imai et al. 2004; Treen et al. 2018). We hypothesized that it could be acting as a maternal repressor of the earliest zygotically expressed genes. To test this, we incubated embryos from the 1-cell stage in high concentrations of FGF, or FGF and the MEK inhibitor U0126 and measured the expression of all the known regulatory genes that are expressed at the 16-cell stage (Satou 2020) by qPCR (Fig S1). We measured samples at the 8-cell stage to see if any genes are precociously expressed by FGF treatment. The only gene that responded to this treatment was *Prdm1-r*.*a*, which encodes a zinc finger transcriptional repressor in *Ciona* (Ikeda et al. 2013). It is linked to *Prdm1-r*.*b*, a paralogous gene that is reported to be expressed from the 32-cell stage and possess overlapping expression and function with *Prdm1-r*.*a* (Ikeda et al. 2013). *Prdm-1r*.*a* was upregulated by FGF treatment at the 8-cell stage, this upregulation could be overcome by adding U0126 (Fig S1).

To verify our qPCR results we developed a 4-color hybridization chain reaction (HCR) *in situ* detection method. We used probes for both *Prdm1-r*.*a, Prdm1-r*.*b*, the known FGF responsive gene *Otx* (Bertrand et al. 2003), as well as the earliest expressed zygotic gene, *Foxa*.*a* (Fig 1A). For each gene, 2 large spots could be observed within the nucleus, which are likely to correspond to sites of active transcription. This observation was supported by the colocalization of the large spots for *Prdm1-r*.*a* and *Prdm1-r*.*b*, as the transcriptional start sites for these genes are only approximately 18 kb apart (Satou et al. 2019). Smaller spots could also be detected within the nucleus as well as in the cytoplasm that are presumably clusters of RNAs, or possibly individual transcripts. 4-color HCR *in situ* was able to recapitulate previously described gene expression patterns from the 8- to 32- cell stages (Fig 1, S2-S3, Lamy et al., 2006; Ikeda et al., 2013; Williaume et al., 2021) as well as reliably distinguish between cells that are actively transcribing genes, from those cells where the HCR *in situ* signal is exclusively cytoplasmic due to expression at earlier stages.

**Fig.1.**
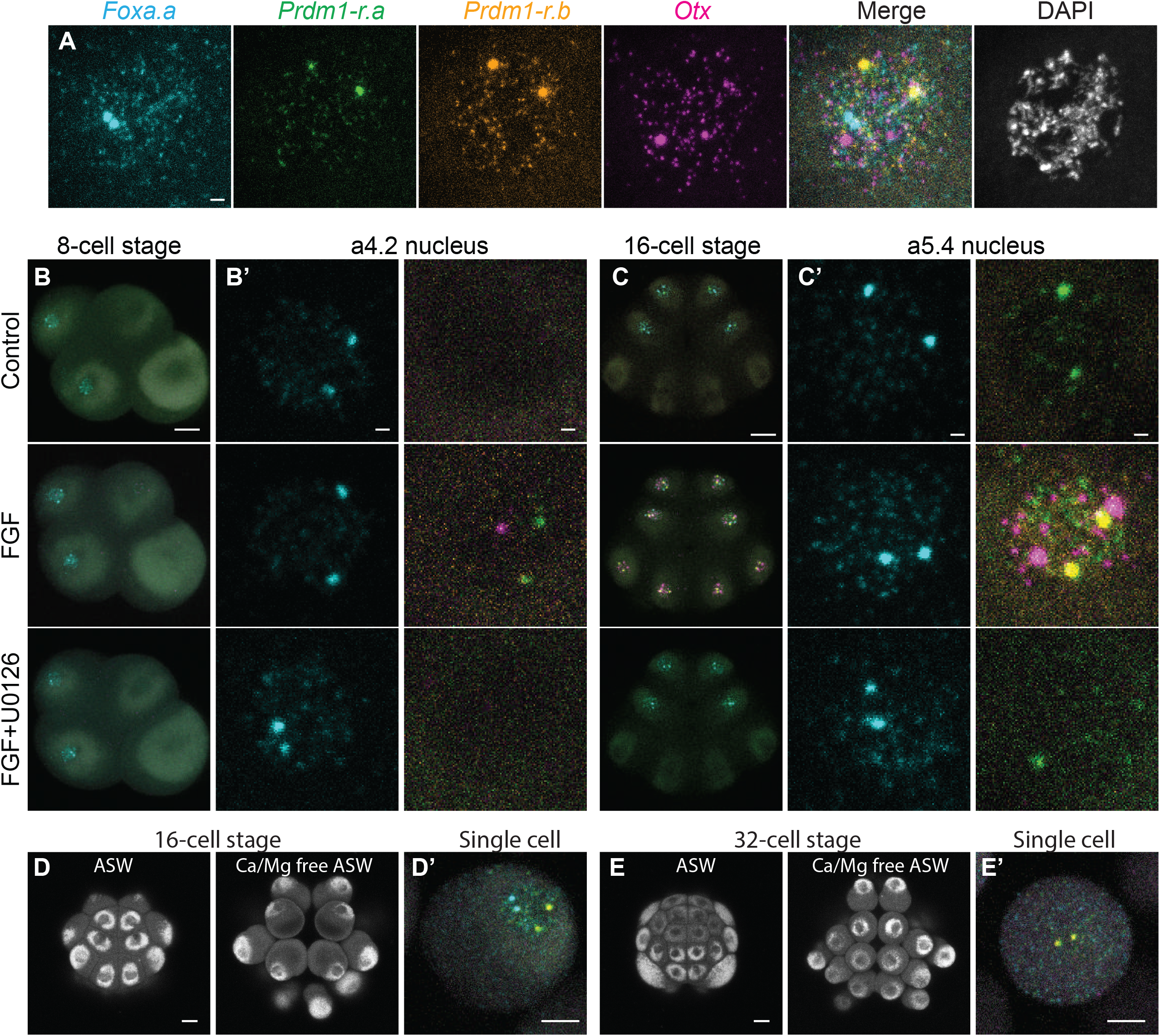
FGF/ERK signaling can induce *Prdm1-r*.*a/b* and *Otx* expression at the 8/16-cell stages. (A) 4-color multiplex HCR fluorescence *in situ* hybridization can detect the expression of *Foxa*.*a* (cyan), *Prdm1-r*.*a* (green), *Prdm1-r*.*b* (orange), and *Otx* (magenta). The image shown is a single nucleus from a 16-cell stage *Ciona* embryo treated with FGF to induce the expression of FGF responsive genes. DNA is stained with DAPI (white). (B,C) *In situ* hybridization at the 8-cell stage (B) and 32-cell stage (C) for the genes in (A). Embryos were treated with FGF or with FGF and U0126. Whole embryo and zoomed in views of an a4.2 (B) and a5.4 nucleus are shown. (D,E) Autofluorescence of an intact 16-cell stage (D) and 32-cell stage (E) *Ciona* embryo cultured in artificial sea water (ASW) or in calcium/magnesium free artificial (Ca^2+^/Mg^2+^ free ASW) sea water to dissociate the blastomeres. (D’) *In situ* hybridization of a single dissociated blastomere from a 16-cell stage embryo that was treated with Ca^2+^/Mg^2+^ free ASW from the 8-cell stage. Nuclear expression of *Foxa*.*a* and *Prdm1-r*.*a* can be seen. (E’) *In situ* hybridization of a single dissociated blastomere from a 32-cell stage embryo that was treated with Ca^2+^/Mg^2+^ free ASW from the 16-cell stage. Cytoplasmic expression of *Foxa*.*a* and nuclear expression of *Prdm1-r*.*a/b* can be seen, but no expression of *Otx* is detected. 8-cell stage embryos are oriented lateral view, anterior left, animal hemisphere up. 16-cell embryos are oriented animal view, anterior up. Scale bars = 20 µm for whole embryo, 2 µm for single nucleus and 10 µm for dissociated cell views.

When 8-cell embryos were treated with FGF we could detect extremely weak nuclear HCR *in situ* signals for *Prdm1-r*.*a* in a minority of embryos (Fig 1B). A single example of precocious activation of *Otx* at the 8-cell stage was also observed. Expression of *Prdm1-r*.*a* could not be detected when embryos were treated with both FGF and U0126, consistent with our qPCR results. Overall, the response of the 8-cell embryo to FGF treatment was extremely modest. This contrasted dramatically with the 16-cell stage. Substantial upregulation of *Prdm1-r*.*a* could be seen as well as ectopic and precocious activation of *Prdm1-r*.*a, Prdm1-r*.*b*, and *Otx* in all cells of the 16-cell stage embryo except the transcriptionally silenced B5.2 cells (Fig 1C, Fig S2,S3; Shirae-Kurabayashi et al., 2011). These effects were overcome when embryos were treated with both FGF and U0126. Under these conditions, *Prdm1-r*.*a* nuclear signals could still be detected in a5.3/a5.4 blastomeres, however, the signal appeared weaker than normal (Fig 1C’).

Early embryos tend to be made up of a small number of large cells, making the establishment of diffusible morphogen gradients difficult. In *Ciona* it has been shown that *Otx* expression is dependent on juxtracrine signaling utilizing extensive cell-cell contacts (Ohta and Satou 2013; Williaume et al., 2021), and it is likely that most signaling events in the embryo work this way (Guignard et al., 2020). We tested if *Prdm1-r*.*a* expression was dependent on cell-cell contacts by transferring 8-cell embryos to Ca^2+^/Mg^2+^ free artificial sea water and allowing them to develop until the late 16-cell stage. This treatment eliminated cell-cell contacts and partially, or completely dissociated the embryos (Fig 1D). HCR *in situs* were performed on these dissociated embryos and *Prdm1-r*.*a* was expressed in similar proportions of cells as intact embryos (Fig S4A). By contrast, as expected, *Otx* expression was lost from cells that were dissociated from the 16- to 32- cell stages (Fig 1E, S4B). We conclude that unlike for *Otx*, cell-cell contacts are not required for *Prdm1-r*.*a* expression.

Using *Ciona* embryos, we can precisely control the timing of fertilization and synchronize development. We performed an in depth-analysis of gene expression from the end of the 8-cell stage (a4.2 cells) to the 32-cell stage (a6.5 and a6.6 cells) (Fig 2A, S5) using samples taken in 5-minute intervals. This revealed a staged onset of gene expression within the 16-cell stage. First *Foxa*.*a* is expressed immediately after mitosis, followed by *Prdm1-r*.*a*. Unexpectedly, we detected expression of *Prdm1-r*.*b* at low levels late in the 16-cell stable. This expression was probably too low to be detected with less sensitive methods (Ikeda et al., 2013). At the 32-cell stage *Prdm1-r*.*a, Prdm1-r*.*b*, and *Otx* all came on simultaneously in the a6.5 cells (Fig 2B). In the a6.6 cells the same temporal expression of *Prdm1-r*.*a* and *Prdm1-r*.*b* could be seen, but there was no expression of *Otx* (Fig S5A).

**Fig.2.**
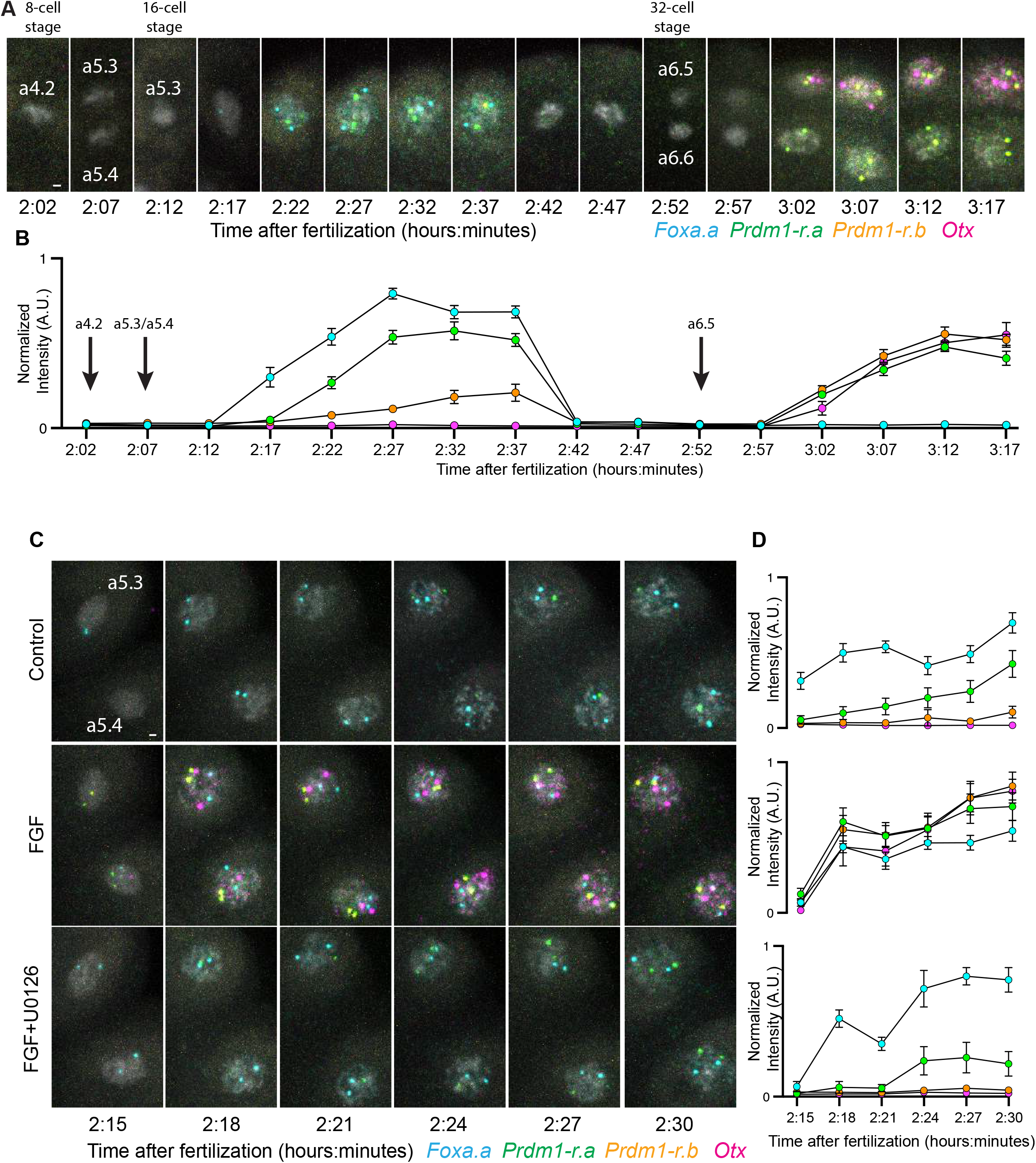
The temporal onset of early zygotic genes. (A) *In situ* hybridization from the 8- to 32- cell stage tracing the a4.2 lineage. DNA is stained with DAPI and shown in white. (B) Quantifications of gene expression levels from samples shown in (A). Each data point indicates the mean (n=24 from 3 embryos) and error bars indicate 95% confidence intervals. (C) *In situ* hybridization from the early to late 16-cell stage. The a5.3 and a5.4 from one half embryo is shown. DNA is stained with DAPI and shown in white. Embryos are treated with FGF or FGF and U0126. (D) Quantifications of gene expression levels from samples shown in (C). Each data point indicates the mean (n=24 from 3 embryos) and error bars indicate 95% confidence intervals. Scale bars = 2 µm

We next investigated how this precise onset of transcription was affected by perturbing FGF signaling. We augmented temporal resolution by using samples taken every 3 minutes. Gene expression was measured during the 16-cell stage in normal development and upon perturbing FGF signaling (Fig 2C,D). When embryos were treated with FGF, we observed *Prdm1-r*.*a, Prdm1-r*.*b*, and *Otx* all expressing simultaneously, immediately after mitosis, at the same time as *Foxa*.*a*. This precocious expression was eliminated when embryos were treated with both FGF and U0126. The kinetics of *Prdm1-r*.*a, Prdm1-r*.*b*, and *Otx* in response to FGF treatment at the 16-cell stage was remarkably similar to what was seen in the a6.5 cell in untreated embryos (Fig 2B,D).

To confirm that our observations are due to repression by maternal ERF, we knocked down translation using morpholino oligonucleotides (MOs). In ERF MO injected embryos faint expression of *Prdm1-r*.*a* (but not *Prdm1-r*.*b* or *Otx*) could be detected at the 8-cell stage (Fig S6). At the 16-cell stage, ERF knockdown resulted in upregulation of *Prdm1-r*.*a, Prdm1-r*.*b*, and *Otx* in all transcriptionally active cells (Fig 3, S7). This demonstrates ERF is a repressor of *Prdm1-r*.*a, Prdm1-r*.*b*, and *Otx* at the 16 cell stage.

**Fig.3.**
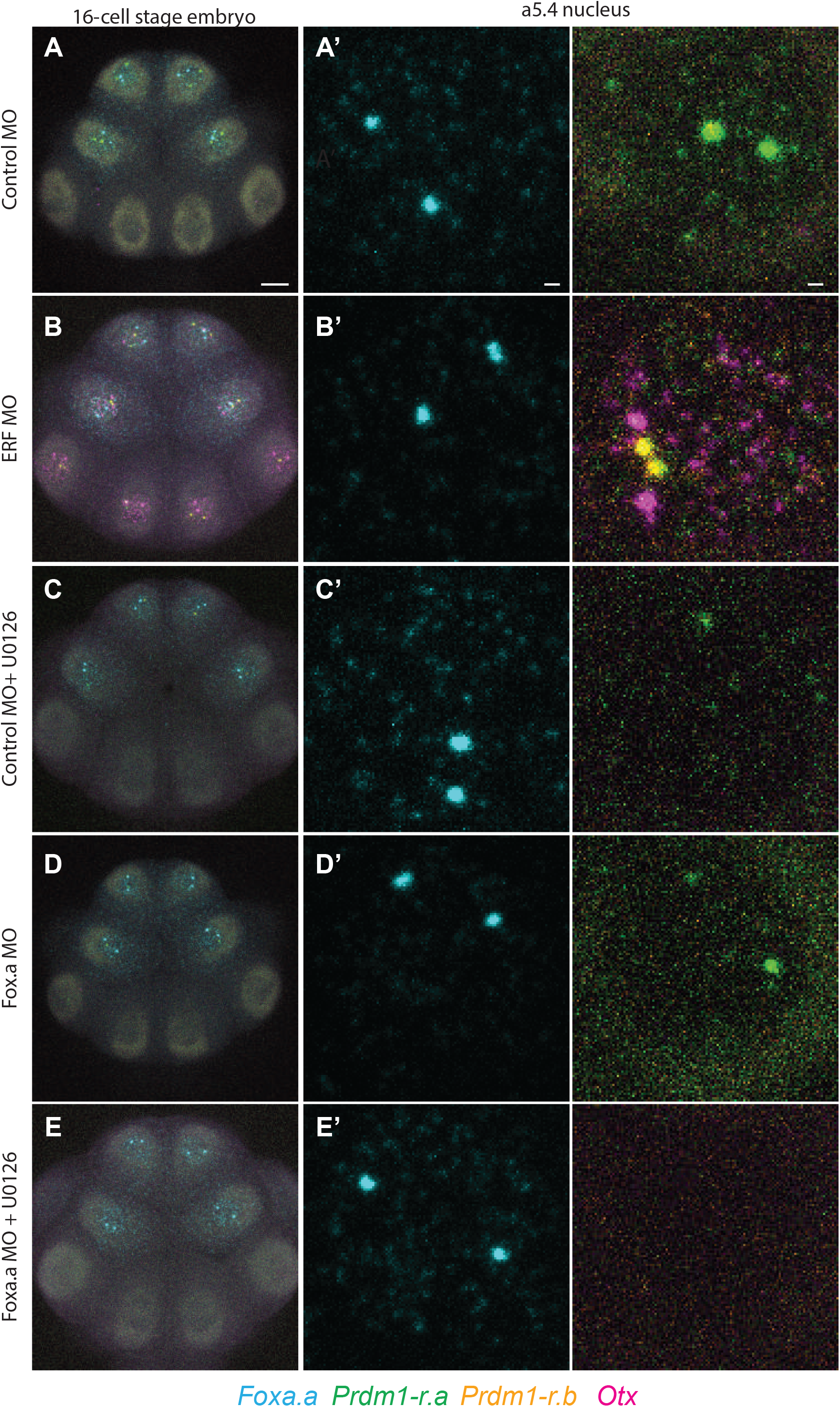
Regulation of *Prdm1-r*.*a/b* and *Otx* by ERF and *Foxa*.*a*. (A) *In situ* hybridizations at the 16-cell stage for embryos injected with a control MO (A’) zoomed in view of the a5.4 nucleus. (B) Same as (A) but embryos are injected with an ERF MO. (C) Same as (A) but embryos are also treated with U0126. (D) Same as (A) but embryos are injected with an Foxa.a MO. (E) Same as (D) but embryos are also treated with U0126. Scale bars = 20 µm for whole embryo and 2 µm for single nucleus views.

Restricted expression of *Prdm1-r*.*a/b* throughout the anterior animal portion of the embryo suggests that *Foxa*.*a* might serve as their activator at the 16-cell stage. To test this, we knocked down *Foxa*.*a by* MO injection. *Prdm1-r*.*a/b* expression is quantitatively reduced in morphants, although weak spots of active transcription could still be reliably detected (Fig 3, S7). Only when we knocked down *Foxa*.*a* and treated embryos with U0126 could we completely abolish *Prdm1-r*.*a* expression at the 16-cell stage (Fig 3, S7), similar to observations at the 32- cell stage (Ikeda and Satou, 2017).

Over the past few years, single cell sequencing atlases have provided comprehensive overviews of gene expression during zygotic genome activation (Xue et al. 2013; Tintori et al. 2016; Treen et al. 2018; Alda-Catalinas et al. 2020; Asami et al. 2022). We also have a general understanding of how early signals can influence what genes are expressed in specific embryonic territories (Stathopoulos et al. 2002; Ohta and Satou 2013; Gentsch et al. 2019).

However, breaking down the precise timing of expression with minute-to-minute accuracy is currently beyond the sensitivity of single-cell RNA seq. In the present study, we employed a combination of 4-color HCR in situ detection methods, along with genetic and experimental perturbation methods to obtain a complete understanding of how a set of signaling responsive genes are activated in early *Ciona* development.

Our results suggest that the embryo is rapidly changing from the 8- to 32- cell stages. At the 8-cell stage, when the embryo is first able to perform zygotic transcription, there appears to be an almost total incompetence to respond to FGF signaling. This can be partially overcome with FGF treatment or ERF knockdown, nevertheless the effects are minimal. At the 16-cell stage, only 30 minutes later, the embryo is fully able to respond to FGF treatment or ERF knockdown by ectopically and precociously expressing FGF responsive genes. The difference between responses at the 8- and 16- cell stages suggest the occurrence of an important transition. While the nature of this switch is unknown, it is easy to imagine that it involves the synthesis or activation of FGF receptors. Because *Prdm1-r*.*a* is still expressed in dissociated embryos, the mechanism does not depend on cell-cell contacts. Instead, autocrine signaling appears to act as an internal timer at the 16-cell stage to delay the onset of *Prdm1-r*.*a/b* expression during the cell cycle (Fig 4A).

**Fig.4.**
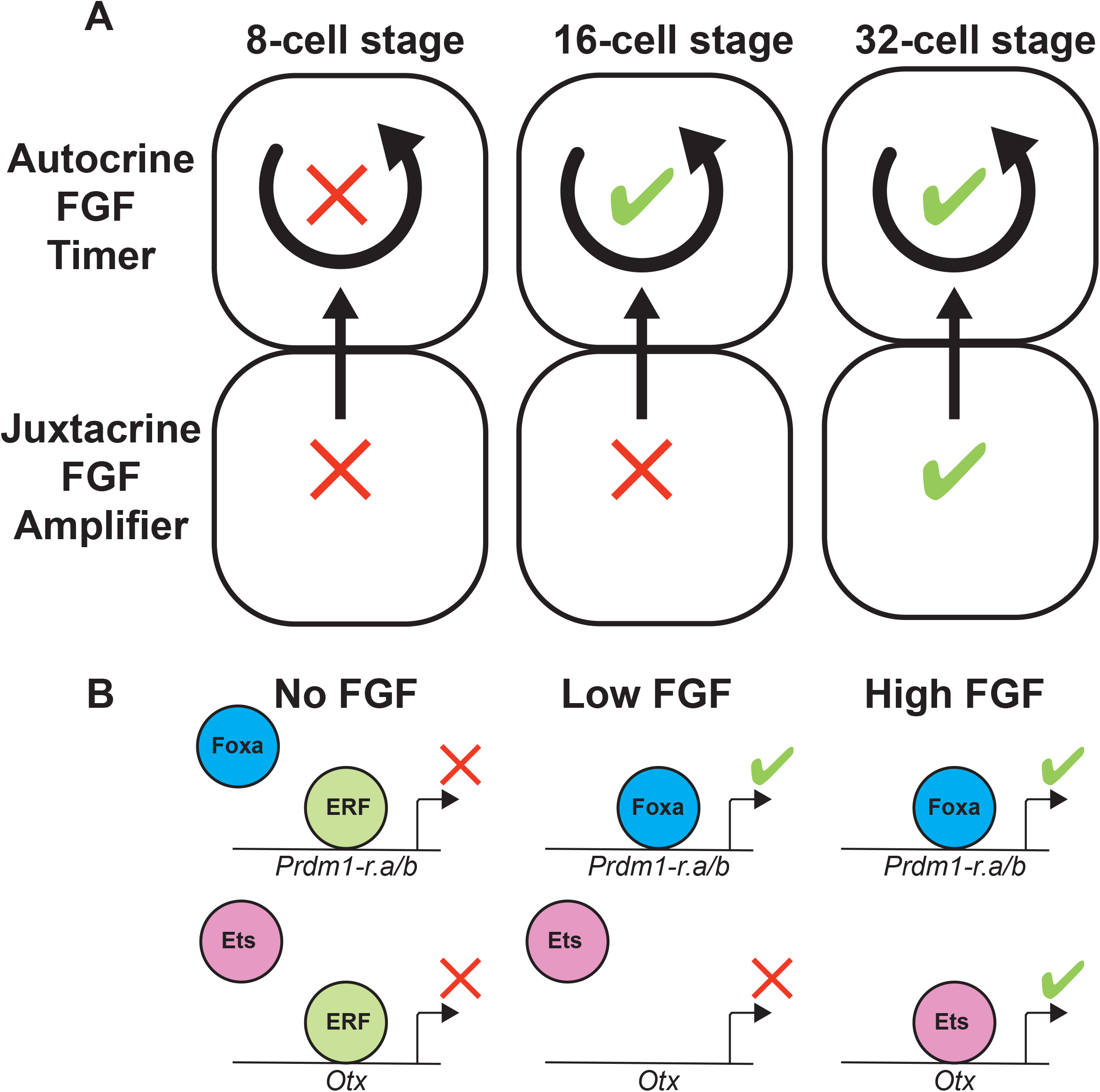
An FGF timer for zygotic genome activation. (A) Schematic depicting an absolute block on FGF signaling at the 8-cell stage that is relieved during the 16-cell stage. At the 32-cell stage, zygotic expression of FGF, as well as compaction of the embryonic cells, has reached a level where it can amplify the signal level. (B) Schematic depicting how the timing mechanisms shown in (A) is interpreted within the nucleus resulting in the repression or activation of *Prdm1-r*.*a/b* and *Otx* at particular developmental stages in response to no, low, or high FGF signals.

Our results suggest that FGF-mediated derepression of ERF allows Foxa.a to activate *Prdm1-r*.*a* and *Prdm1-r*.*b* during the 16-cell stage. However, these low levels of FGF signaling are not sufficient to activate Otx, which might depend on both derepression of ERF and induction of additional activators such as Ets1/2. We propose that temporal precision is tuned during zygotic genome activation by this interplay of localized activators and inactivation of repressors by signaling.

Zebrafish embryos have maternal ERF mRNA at high levels (White et al., 2017). In mouse embryos, ERF does not appear to be maternally expressed, but can be detected in early epiblast stages (Vega-Sendino et al., 2021). We propose that mechanisms similar to those described in *Ciona* are used by a range of vertebrate embryos to control the precise timings of gene expression during zygotic genome activation and early embryogenesis.

## Methods

### Animals

Adult *Ciona intestinalis* Type A (Pacific populations, also referred to as *Ciona robusta*) were sourced commercially from M-REP (San Marco, CA, USA). Live adults and embryos were handled at 18°C.

### Quantitative PCR

3 batches of embryos from different adult individuals were developed until the 8-cell stage. Total RNA purification, cDNA synthesis, and qPCR amplifications and quantifications were done as previously described (Treen et al., 2018). Primers used for qPCR are listed in Table S1.

### Treatments

U0126 (U120, Sigma-Aldrich, St. Louis, MO, USA) stock solutions were dissolved in DMSO and embryo treatment concentrations were 10 μM. FGF treatments were done using recombinant Fibroblast Growth Factor-Basic (F3685, Sigma-Aldrich) concentrations were 300 ng/mL. Control samples were treated with 0.1% DMSO. All treatments were done at the 1-cell stage approximately 30 minutes after fertilization.

### Microinjections

Antisense morpholino oligonucleotides (MOs) were commercially synthesized by Gene Tools (Philomath, OR, USA). The MO sequences were: *ERF* 5’- CACATACGAGCAGTGCATGATTAAG 3’ *Foxa*.*a* 5’-GAGACGACAACATCATTTTTGTAC 3’.

Control MOs used the Gene Tools standard control oligo (5’- CCTCTTACCTCAGTTACAATTTATA 3’). MO injection concentrations were 0.5mM Microinjections were done as previously described (Treen et al., 2018).

### Hybridization chain reaction probe design

Probes sets for HCR in situ hybridization were produced commercially by Molecular Instruments (Los Angeles, CA, USA) based on the following gene models: KY21.Chr11.1129.v1.SL1-1 (*Foxa*.*a*), KY21.Chr12.997.v1.SL1-1 (*Prdm1-r*.*a*), KY21.Chr12.994.v1.SL1-1 (*Prdm1-r*.*b*), KY21.Chr4.720.v2.SL2-1 (*Otx*). To ensure no cross hybridization between *Prdm1-r*.*a* and *Prdm1-r*.*b* probe sets, the probe sets were designed to target nucleotides 1-844 (*Prdm1-r*.*a*) and 1-877 (*Prdm1-r*.*b*) of the predicted transcripts. Probes were designed to be compatible with the following HCR amplifiers: Foxa.a – B2, Prdm1-r.a – B1, Prdm1-r.b – B5, Otx – B3. 20 split initiator pairs were used for each gene.

### Hybridization chain reaction *in situ* hybridization

Embryos were fixed in 100 mM HEPES, 500 mM NaCl, 2 mM MgSO_4_, 2 mM EGS (ethylene glycol bis(succinimidyl succinate)), 1% formaldehyde for 5 minutes with constant agitation and then for a further 55 minutes without agitation at room temperature. Samples were then washed in phosphate buffered saline, 0.1% Tween 20 (PBST) 4 times. Samples were then dehydrated by replacing the PBST with 50% ethanol 2 times and then 80% ethanol 2 times. Samples were stored in 80% ethanol at −20°C for between 1-7 days before use.

The HCR protocol was based on a previously published protocol for sea urchin embryos (Choi et al, 2018) with some modifications. Dehydrated embryos were rehydrated from 80% ethanol by gradually adding 5x Sodium chloride sodium citrate, 0.1% Tween 20 (5x SSCT) until the remaining ethanol concentration was approximately 20%. Samples were then washed in 5x SSCT 3 times. Rehydrated samples were pre-hybridized in a pre-hybridization buffer (5x Sodium chloride sodium citrate, 50% formamide, 5x Denhardt’s solution, 100 μg/mL yeast tRNA, 100 μg/mL salmon sperm DNA) for 2 hours at 37°C. We found that shorter pre-hybridization times of 30 minutes to 1 hour, as well as the absence of yeast tRNA and salmon sperm DNA in the pre-hybridization solution resulted in non-specific fluorescence signals at cell surfaces. The pre-hybridization buffer was then replaced with 0.8 pmol of each HCR probe set diluted in fresh pre-hybridization buffer and incubated for 16-18 hours at 37°C. Samples were washed in HCR wash buffer (Molecular Instruments) for 5 minutes 2 times and then for 30 minutes 2 times at 37°C. Samples were washed twice in 5x SSCT at room temperature. The amplification solution was prepared by heating and snap cooling HCR hairpins: Alexa 488-B1, Alexa 647 – B2, Alexa 514 – B5, Alexa 546 -B3 as previously described (Choi et al 2018). 6 pmol of each HCR hairpin was diluted in HCR amplification buffer (Molecular Instruments).

Samples were incubated with HCR hairpins diluted in HCR amplification buffer for 3 hours at room temperature in the dark. Samples were washed in 5x SSCT for 5 minutes 2 times and then for 30 minutes 2 times. In the penultimate wash, DAPI was added to stain DNA. The 5X SSCT was replaced with PBST and samples were kept in the dark at 4°C until they were imaged.

### Imaging

HCR *in situ* samples were imaged using a Zeiss LSM 880 confocal microscope (Carl Zeiss, Oberkochen, Germany) using a 20x/0.8NA plan-apochromat objective. Z-section of whole embryos were taken with a pixel size of 0.149 μm^2^ and 1 μm Z-stack steps. In order to minimize cross talk each fluorophore imaged separately (except DAPI and Alexa 647, which were imaged simultaneously) was using the following conditions: Alexa 488, 488 nm laser, 488-533 nm emission filters; Alexa 514, 514 nm laser, 544-554 nm emission filters; Alexa 546, 561 nm laser, 597-633 nm emission filters; DAPI/Alexa 647, 405 and 633 nm lasers, 415-480 and 649-733 nm emission filters.

### Quantification of HCR in situ signals

For each gene, the mean fluorescence intensity of a 2 μm diameter circular region was measured for the brightest two spots within the nucleus (assumed to be the sites of active transcription) at the Z-section that had the highest signal. If these signals were absent, the background signal of a random region within the nucleus was measured. Fluorescence intensity was normalized for each individual gene within a dataset by dividing each signal level by the highest measurement for that gene giving a data range of 0-1.

## Acknowledgements

We thank members of the Levine lab for helpful discussions. This work was supported by grants from the NIH (NS076542) and Princeton Catalysis Initiative.

## Figure Legends

**Fig.S1.**
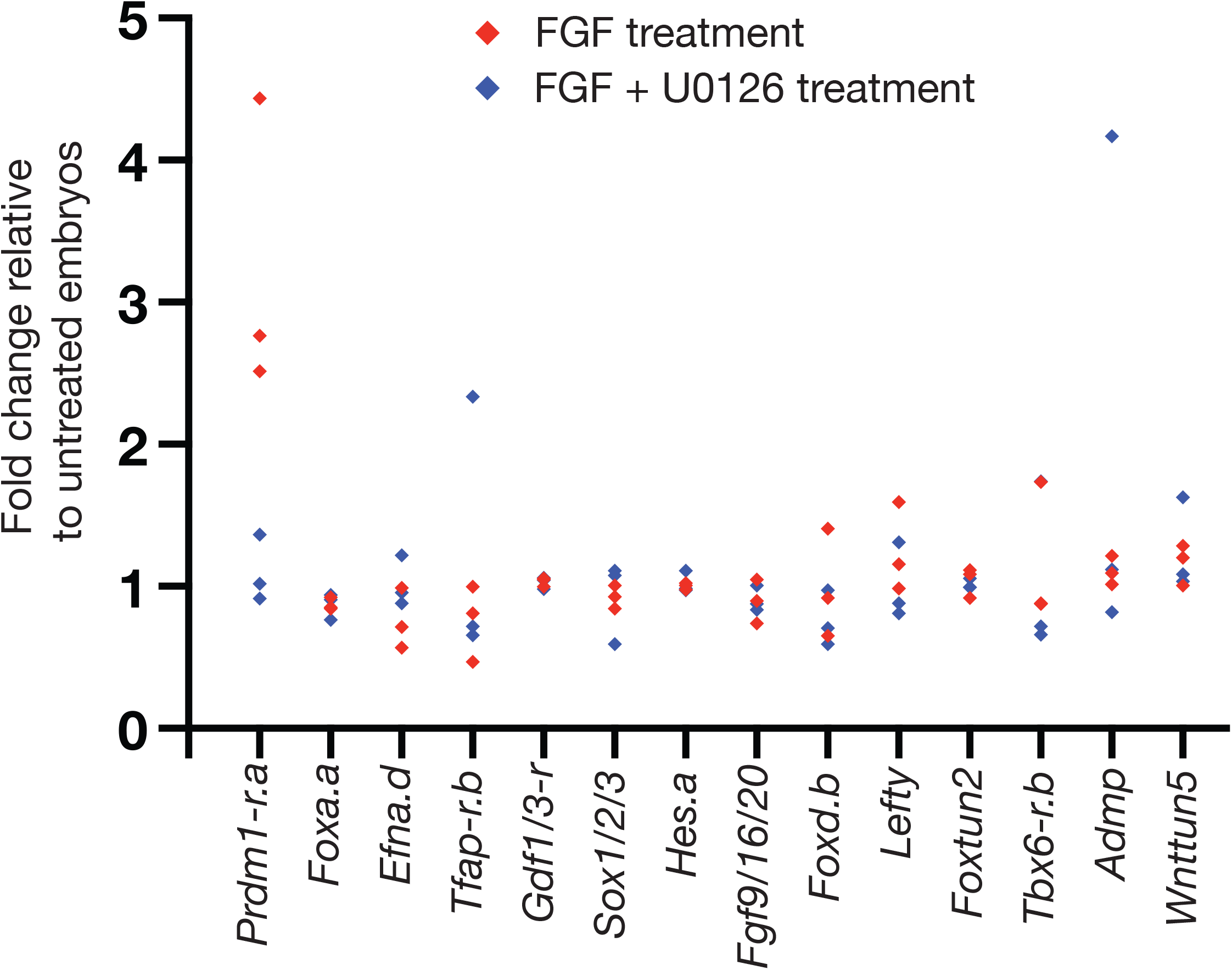
The response of early zygotic genes to FGF treatment. qPCR analysis of the expression levels of all the known regulatory genes that are expressed at the 16-cell stage. Samples were treated with FGF or FGF + U0126 from the 1-cell stage and RNA was isolated at the 8-cell stage. Each data point is measured from several hundred pooled embryos.

**Fig.S2.**
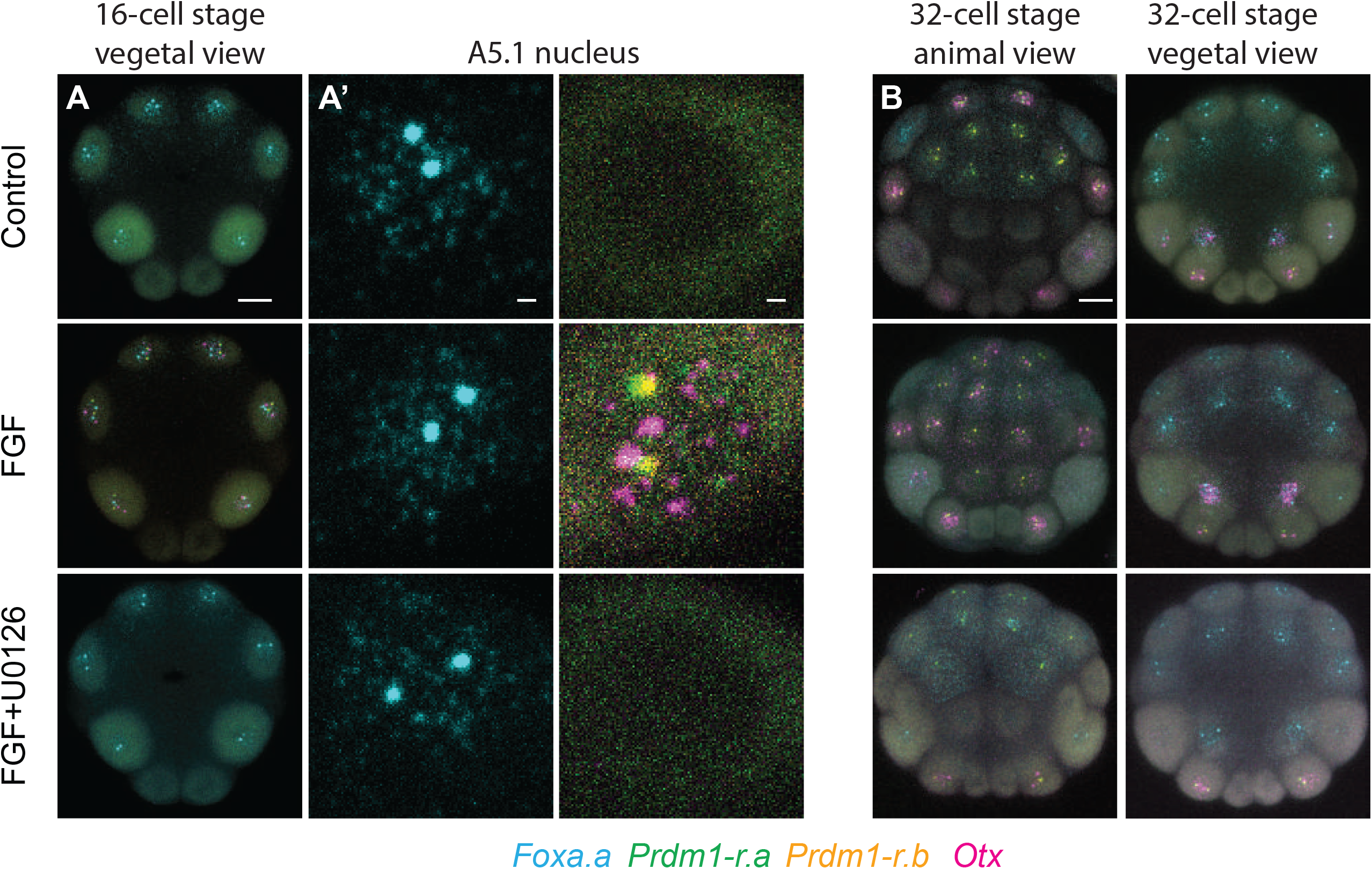
Additional examples of early *Ciona* embryo responses to FGF/ERK signal perturbations. (A) *In situ* hybridization at the 16-cell stage. Embryos were treated with FGF or with FGF and U0126. (A’) zoomed in views of A5.1 nucleus are shown. (B) *In situ* hybridization at the 32-cell stage. Embryos were treated with FGF or with FGF and U0126. 16-cell embryos are oriented vegetal view, anterior up. 32-cell embryos are oriented as indicated in the figure, anterior up. Scale bars = 20 µm for whole embryo and 2 µm for single nucleus views.

**Fig. S3.**
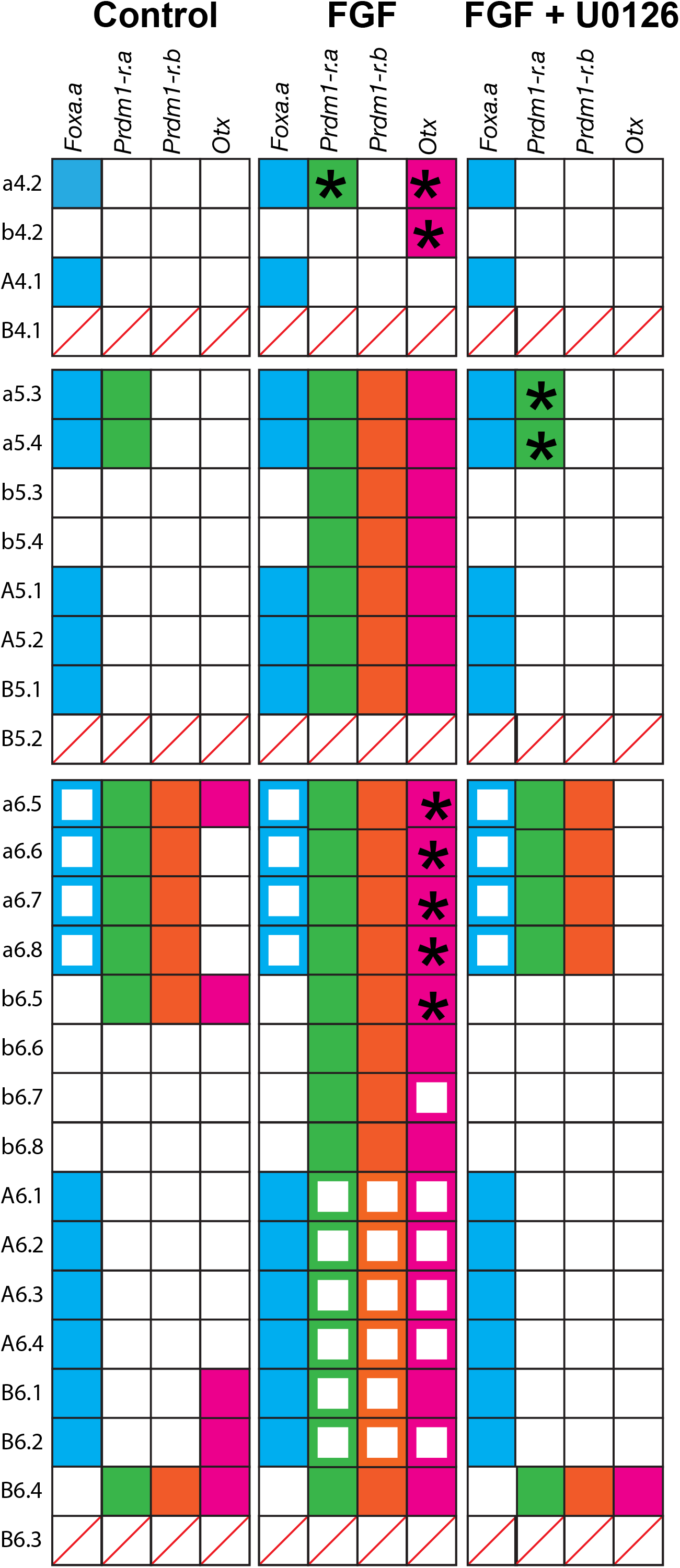
Summary of gene expression patterns observed for FGF and U0126 treatments. If a gene is expressed in a specific cell at a specific time point this is indicated be a colored box. Cytoplasmic expression (from the previous stage) is indicated by a hollow colored box. Instances where expression is considerably lower than controls or is only detected in a small subset of embryos or vary considerably from batches are indicated with an asterisk.

**Fig. S4.**
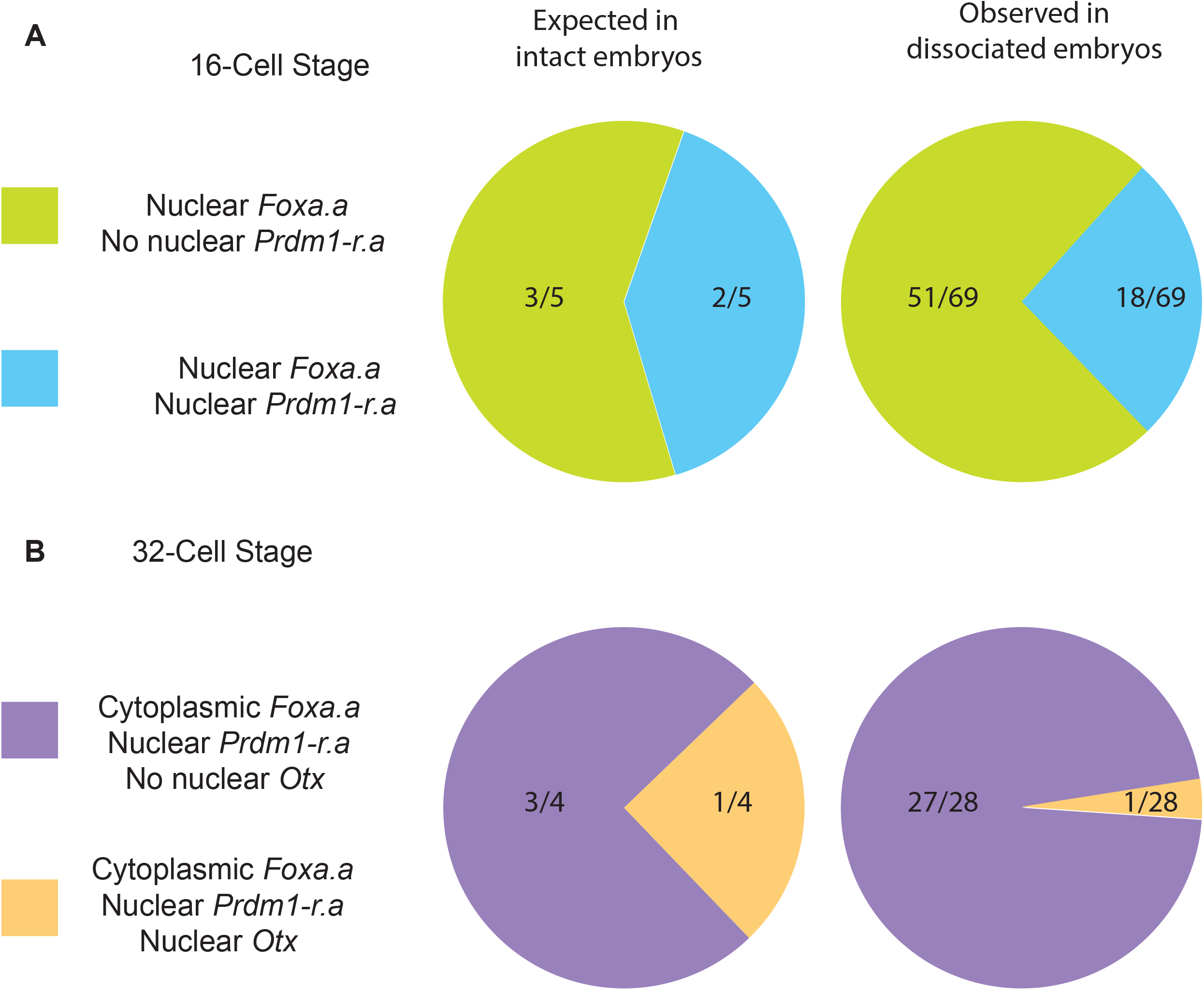
Gene expression in dissociated cells. (A) Pie chart showing the numbers of cells expected to have the indicated genes expressed or not expressed in intact embryos, or observed in dissociated embryos at the 16-cell stage. (B) same as (A) but for the 32-cell stage.

**Fig. S5.**
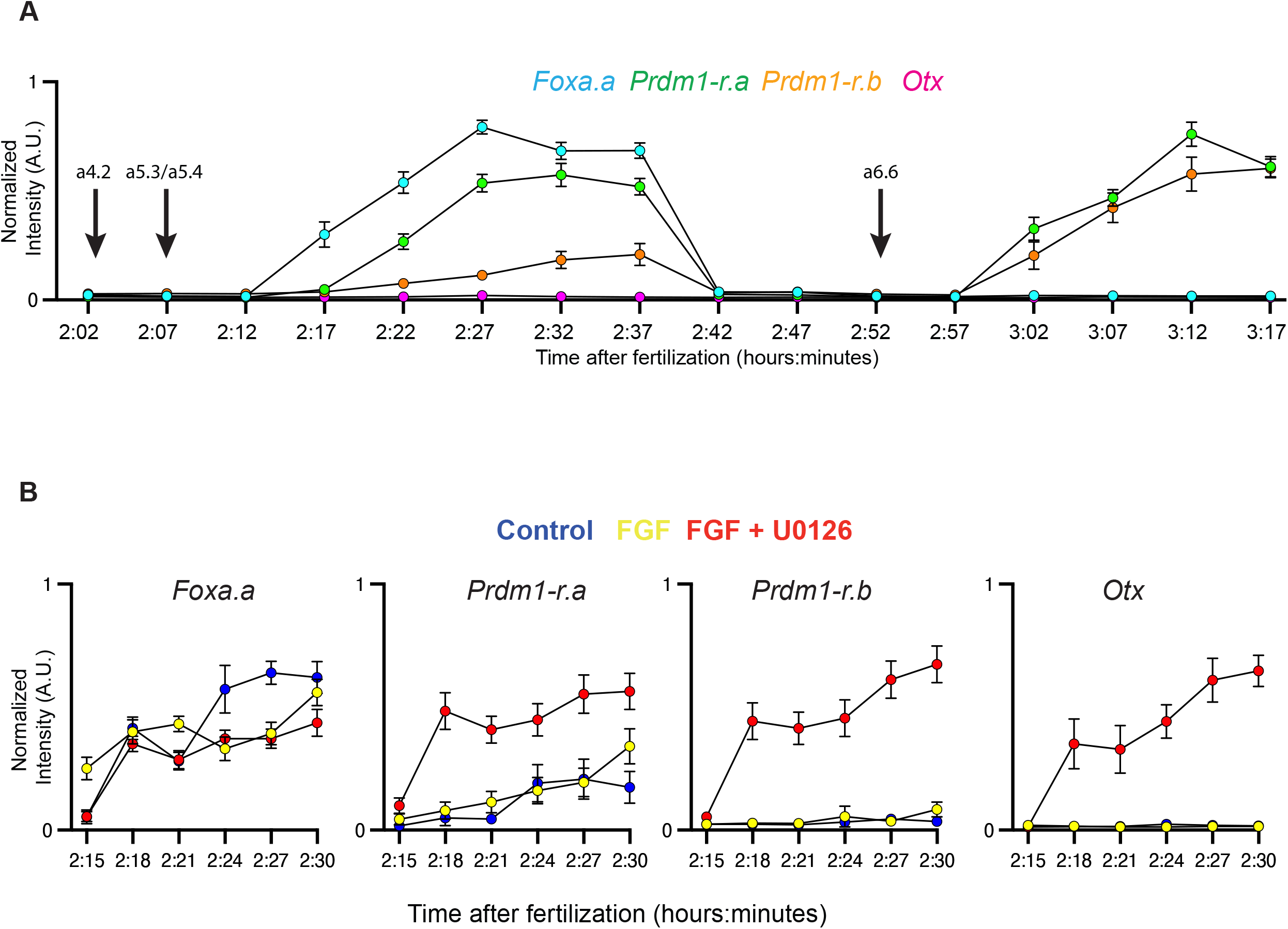
Additional quantifications of temporal onset of early zygotic genes. (A) Graph depicts the same dataset as Fig. 2B but measures expression in the a6.6 nucleus at the 32-cell stage. (B) Graphs depict the same dataset in Fig. 2D but are sorted by individual genes.

**Fig. S6.**
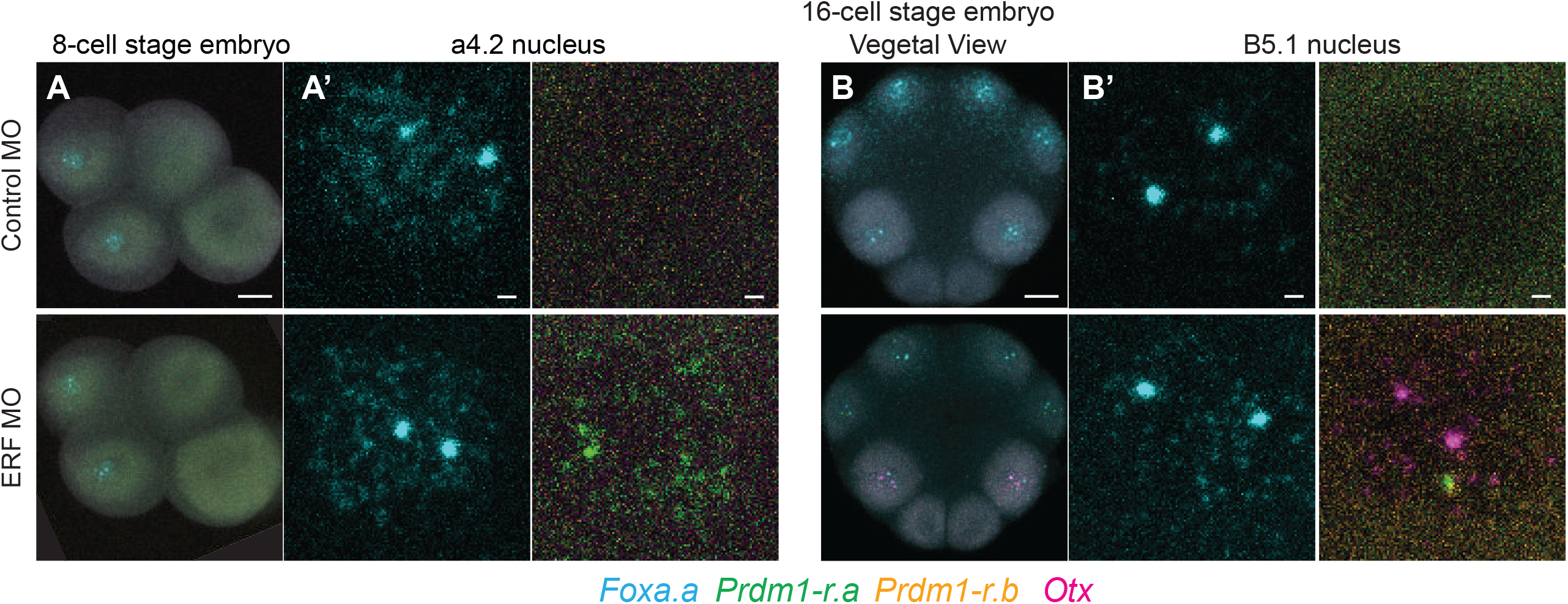
Additional examples of regulation of *Prdm1-r*.*a/b* and *Otx* by ERF. (A) *In situ* hybridizations at the 16-cell stage for embryos injected with control MO or ERF MO (A’) zoomed in view of the a5.4 nucleus. (B) Same as (A) but for the 16-cell stage. 8-cell stage embryos are oriented lateral view, anterior left, animal hemisphere up. 16-cell embryos are oriented vegetal view, anterior up. Scale bars = 20 µm for whole embryo and 2 µm for single nucleus views.

**Fig. S7.**
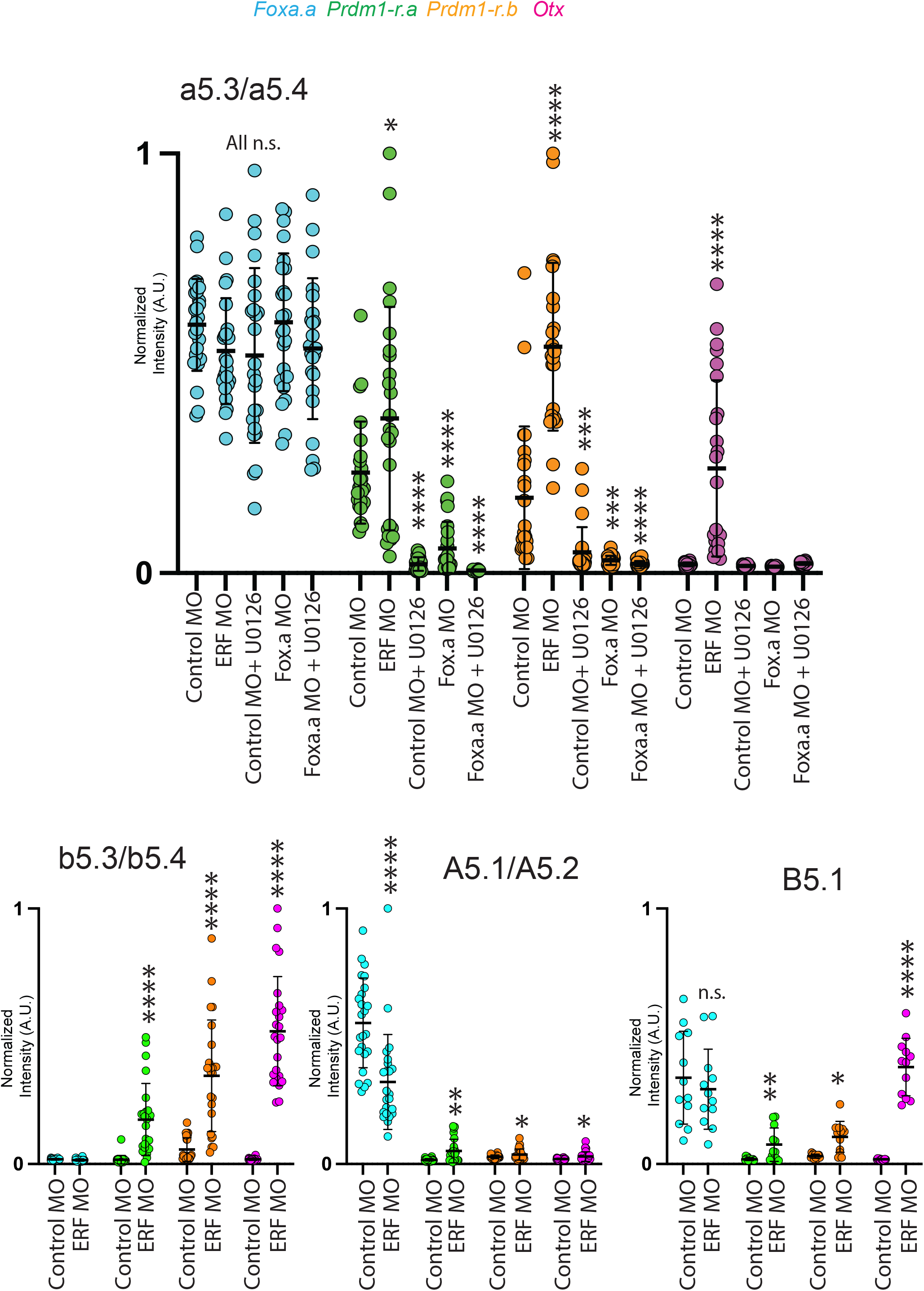
Quantifications of knockdown experiments. Graphs show quantifications of MO knockdown experiments performed in this study. Each data point is the normalized fluorescence of a single site of active transcription. Error bars depict mean and standard deviations. Statistical significance was determined using paired two-tailed t tests. n.s. = not significant, *= p value <0.05, **= p value <0.01, ***= p value <0.001, ****= p value <0.0001.

**Table S1.**
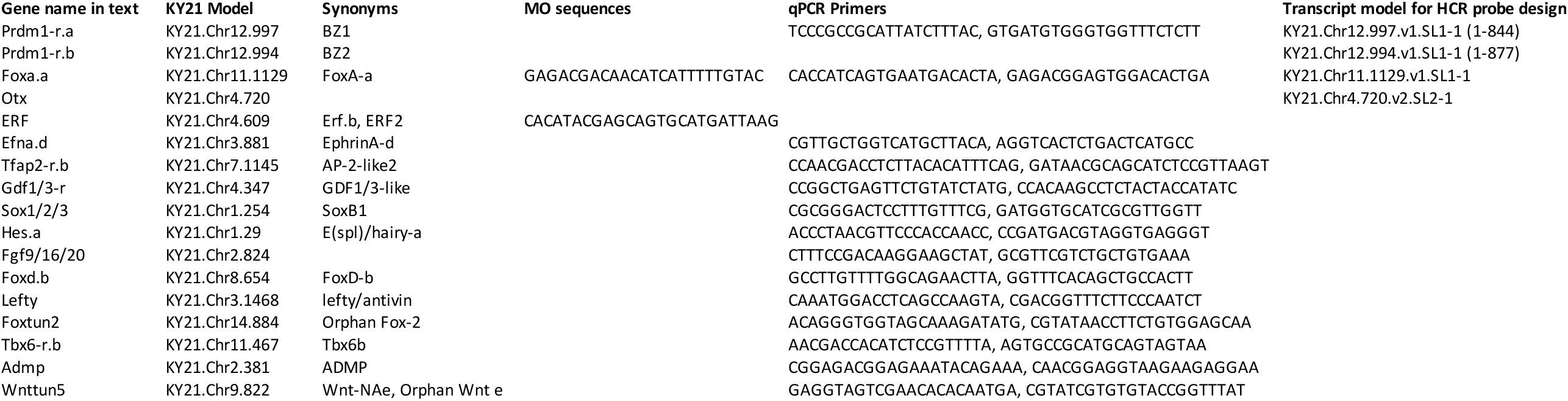

